# Unravelling a mechanistic link between mitophagy defect, mitochondrial malfunction, and apoptotic neurodegeneration in MPS VII

**DOI:** 10.1101/2024.08.23.609363

**Authors:** Nishan Mandal, Apurba Das, Rupak Datta

## Abstract

Progressive neurodegeneration and cognitive disability are prominent symptoms of MPS VII, a lysosomal storage disorder caused by β-glucuronidase enzyme deficiency. Yet, the mechanism of neurodegeneration in MPS VII remains elusive thereby limiting the scope of targeted intervention. To address this, we recently developed a β-glucuronidase-deficient (CG2135^-/-^) *Drosophila* model of MPS VII. The CG2135^-/-^ flies exhibited signs of neuromuscular degeneration, including loss of dopaminergic neurons, and accumulated engorged lysosomes, ubiquitinated proteins and mitochondria in their brains. These observations, coupled with our current finding that the CG2135^-/-^ flies were highly susceptible to starvation, prompted us to investigate potential defects in the autophagy-lysosomal clearance machinery in the brain. We found that both autophagy induction and lysosome-mediated autophagosomal turnover were impaired in the CG2135^-/-^ fly brain. This was evidenced by lower Atg8a-II levels, reduced Atg1 and Ref(2)P expression along with accumulation of lipofuscin-like inclusions and multilamellar bodies. Interestingly, mitophagy was also found to be defective in their brain, causing buildup of enlarged mitochondria with distorted cristae and reduced membrane potential. This, in turn, affected mitochondrial function as reflected by drastically reduced brain ATP levels. Energy depletion triggered apoptosis in neuronal as well as non-neuronal cells of the CG2135^-/-^ fly brain, where we also detected apoptotic dopaminergic neurons. Resveratrol treatment, previously found to protect against loss of dopaminergic neurons in the CG2135^-/-^ flies, has now been shown act by correcting the mitophagy defect and preventing ATP depletion. Collectively, our study establishes a causal link between mitophagy defect, mitochondrial malfunction, and apoptotic neurodegeneration in MPS VII.

## Introduction

Mucopolysaccharidosis VII (MPS VII) is a recessively inherited lysosomal storage disorder caused by mutations in the *β-glucuronidase* (*β-GUS*) gene. β-GUS is one of the eleven glycosaminoglycan (GAG)-catabolizing lysosomal hydrolase, deficiency of which results in accumulation of undegraded or partially degraded GAGs in the lysosomes ^1,2^. MPS VII patients suffer from progressive multiorgan dysfunction, and most of them eventually succumb to premature death. Common clinical manifestations of MPS VII include, hydrops fetalis, short stature with coarse face and skeletal deformities, heart valve abnormality, hepato-splenomegaly, and recurrent infections ^1–3^. Additionally, MPS VII patients also develop a variety of neurological symptoms like mental retardation, limited vocabulary, delayed speech, hearing impairment and restricted movement ^3,4^. This indicates that MPS VII patients suffer from neurodegeneration, which was corroborated by autopsy studies showing extensive lysosomal storage in different regions of the patient’s brain ^5,6^. The MPS VII mouse model also exhibited accumulation of storage vacuoles throughout the brain and signs of neurodegeneration in the hippocampus and cerebral cortex regions that worsened with age ^7^. However, the molecular and cellular events that connect lysosomal storage to neurodegeneration in MPS VII are still not well-understood. Without this knowledge, developing a mechanism-based therapy to ameliorate neurological problems of MPS VII patients remains an unmet need. This issue has become even more critical because enzyme replacement therapy (ERT), the only approved treatment for MPS VII patients, despite being successful in alleviating some of the disease symptoms, has not yet been shown to resolve the neurological complications ^8,9^. This is most likely due to the limited ability of the infused enzyme to cross the blood brain barrier ^9^. All these challenges emphasize the urgent need for an in-depth study of the brain pathology and the mechanism of neurodegeneration in MPS VII that would facilitate exploration of small molecule-based therapy.

Towards this goal, recently we developed the first *Drosophila* model of MPS VII by knocking-out the fly *β-GUS* gene (*CG2135*) ^10^. The CG2135^-/-^ fly exhibited progressive locomotor disability, attributed to neuromuscular degeneration. The CG2135^-/-^ fly brain displayed distinctive features like, extensive vacuolation, abnormal buildup of engorged lysosome as well as ubiquitinated proteins and an increased abundance of MitoTracker-positive vesicles. Interestingly, a significant loss of dopaminergic neurons was also observed in the CG2135^-/-^ fly brain ^10^. Dopaminergic neurons are classically known to control locomotion and their depletion is associated with movement disorders such as Parkinson’s disease ^11,12^. These initial findings provided us with an excellent platform for delving deeper into the underlying mechanism of neurodegeneration in MPS VII.

In line with our observation in the CG2135^-/-^ fly brain, Martins *et. al.* previously reported mitochondrial accumulation in the MPS IIIC mouse brain. Their data also revealed a progressive decline in activities of some of the respiratory chain enzymes in mitochondria isolated from the MPS IIIC mice brain ^13^. Apart from these two MPS disorders, mitochondrial accumulation in brain was also reported in animal models of few other unrelated lysosomal storage disorders, e.g. Gaucher disease, multiple sulfatase deficiency and CLN7 ^14–16^. In this context, it is worth mentioning that, due to their high energy demand, neuronal cells rely heavily on mitochondrial quality control to maintain a healthy and efficient pool of mitochondria. These quality control mechanisms include biogenesis of mitochondria, their fusion/fission dynamics and selective clearance of damaged mitochondria through the mitophagy pathway ^17^. Aberrations in these processes have been reported to play a role in many neurodegenerative diseases ^18,19^. However, very little is known about the mechanistic link between mitochondrial quality control in the brain and neurodegeneration associated with MPS disorders in general, and MPS VII in particular.

In this study, we systematically investigated the causes and consequences of abnormal buildup of mitochondria in the CG2135^-/-^ fly brain to better understand the mechanism of neurodegeneration in MPS VII. We initially focused on the autophagy-lysosomal clearance pathway in the brain to check if any defect in this machinery led to inefficient turnover of mitochondria. This notion was fuelled by our previously reported observation of aberrant autophagy in the CG2135^-/-^ larvae ^20^. A similar defect was also reported in β-GUS deficient chondrocyte cell line ^21^. Autophagy is a coordinated cellular waste disposal system in which cell debris is packed into double-membrane autophagosome vesicles and targeted to lysosome for degradation and recycling ^22^. The process involves a series of autophagy-related genes (Atg) and proteins which are regulated by various physiological cues ^23^. Apart from bulk autophagy, there are specialized forms of autophagy such as ER-phagy or mitophagy, via which damaged cellular organelles are specifically degraded ^19,24^. We first suspected a gross autophagy defect in adult CG2135^-/-^ flies when they were found to be highly susceptible to starvation. Following this lead, we confirmed that autophagy is indeed compromised in the CG2135^-/-^ fly brain. Further probing uncovered that the process of mitophagy was also impaired in the CG2135^-/-^ fly brain, resulting in the accumulation of abnormally large mitochondria with deformed cristae structure and decreased membrane potential. Consequently, ATP levels dropped significantly, triggering extensive apoptotic cell death in the CG2135^-/-^ fly brain, including dopaminergic neurons. Treatment with resveratrol, previously shown to prevent loss of dopaminergic neurons in the CG2135^-/-^ flies, has now been found to protect the CG2135^-/-^ fly brain from mitophagy defect and ATP exhaustion. Our findings thus established a mechanistic link between mitophagy defect, mitochondrial malfunction, and apoptotic neurodegeneration in MPS VII.

## Results

### Defective autophagy-lysosome clearance system in CG2135^-/-^ fly brain

During initial characterization of the MPSVII fly (CG2135^-/-^), we observed an abnormal accumulation of MitoTracker-positive puncta and ubiquitinated proteins in their brain, indicating a potential defect in the lysosome-mediated cellular clearance system ^10^. In a follow-up study, we reported aberrant autophagy in the CG2135^-/-^ larvae ^20^. These findings prompted us to analyze the status of autophagy in the adult CG2135^-/-^ fly, specifically in its brain, to better understand the mechanism of neurodegeneration in this fly model. Autophagy pathway is known to be activated as a protective response to nutrient starvation ^25^. Studies with various model organisms have established that an autophagy defect results in enhanced susceptibility to starvation ^26–28^. Taking cue from these reports, we compared the starvation sensitivity of the adult CG2135^-/-^ flies with their age-matched wild-type counterparts. For this, we took wild type (W^1118^) and CG2135^-/-^ flies of different age groups (young flies - 4 days old, middle-aged flies - 15 days old and old flies - ≥ 30 days old) and subjected them to a no-food condition. The percentage survivability was analyzed every 12 hours by counting the number of surviving flies (Figure 1A). The survival curves, as well as the mean survival time (MST) under starved condition, clearly showed that the CG2135^-/-^ flies were more susceptible to starvation than the wild type flies at all ages (Figures 1B-1D). This data strongly indicated that the adult CG2135^-/-^ flies could be suffering from an autophagy defect. We then shifted our focus to a detailed analysis of the autophagy status in the CG2135^-/-^ fly brain. For this, we collected the wild type and CG2135^-/-^ flies of different age groups (4-days old to 45-days old) and examined the level of Atg8a, the *Drosophila* homolog of mammalian LC3, in their brains. Atg8a is considered as a reliable marker for autophagy as it gets recruited to autophagosome vesicles in its phosphatidylethanolamine (PE) conjugated form ^29^.Our western blot data clearly showed a marked reduction in the Atg8a-II levels (the PE-conjugated form of Atg8a that correlates with the autophagosome abundance) in the CG2135^-/-^ fly brain as compared to their wild type counterparts across all age groups. Notably, the differences in brain Atg8a-II levels between the wild type and the CG2135^-/-^ fly progressively increased with age (Figures 2A and 2B). Consistent with this finding, immunofluorescence staining of whole fly brains with an Atg8a antibody showed a significant reduction in the Atg8a-positive autophagic vesicles in 30-days old CG2135^-/-^ fly brain compared to age-matched control (Figures. S1A-F). In this context, it is worth noting that although there was no significant difference in the Atg8a transcript levels between the wild type and the CG2135^-/-^ fly brains, the Atg8a-I protein level (the non-autophagosome bound form of Atg8a) was also found to be reduced in the CG2135^-/-^ fly brains (Figures S1G – S1I). To ascertain the reason behind the reduced abundance of autophagosomes in the CG2135^-/-^ fly brain, we checked the status of the Ref(2)P, the fly homologue of the mammalian autophagic cargo adapter protein p62/SQSTM1 ^30^. As evident from our whole-brain immunofluorescence staining and western blot data, the Ref(2)P protein level in the CG2135^-/-^ fly brain was significantly less as compared to the wild type flies of similar age (Figures 2C – 2I). To study if lowering of the Ref(2)P protein level is due to transcriptional downregulation, we analyzed the Ref(2)P mRNA level by RT-qPCR and found reduced by ∼ 40% in the CG2135^-/-^ fly brain (Figure 2J). This prompted us to also check the expression of Atg1, one of the important autophagy initiation genes ^31^. Interestingly, almost 60% decline in the transcript level of Atg1 was observed in the CG2135^-/-^ fly brain as compared to their age-matched wild type counterparts (Figure 2K). Collectively, these data indicate that a defect in the autophagy induction might be the underlying reason behind the reduced abundance of the autophagosomes in the old CG2135^-/-^ fly brain. We also noticed that in stark contrast to the wild type flies, the optical lobe and midbrain region of the 45-days old CG2135^-/-^ flies exhibited distinctive autofluorescence when imaged in the FITC channel (Figures 2L – 2N). Such autofluorescent puncta are reminiscent of lipofuscin, the storage granules of undegraded lipids and misfolded proteins that accumulate in the lysosomes of postmitotic neurons ^32^. Accumulation of lipofuscin is a hallmark of ageing and also has been reported in many neurodegenerative diseases ^33–35^. This intriguing finding prompted us to further analyze the tissue architecture of the fly brains using high resolution transmission electron microscopy (TEM). Our TEM images revealed that compared to the wild type flies, the CG2135^-/-^ fly brains contained very few double-membrane autophagosome vesicles but instead had an abundance of multilamellar bodies (MLBs) with whorl-like structures (Figures 2O– 2S). Abnormal accumulation of MLBs has been observed in a variety of lysosomal storage disorders, resulting from dysfunction of lysosomal hydrolytic activity ^34,36,37^. Considering all these findings, it is evident that both autophagy induction and lysosome-mediated autophagosomal clearance were impaired in the CG2135^-/-^ fly brain.

**Figure 1:**
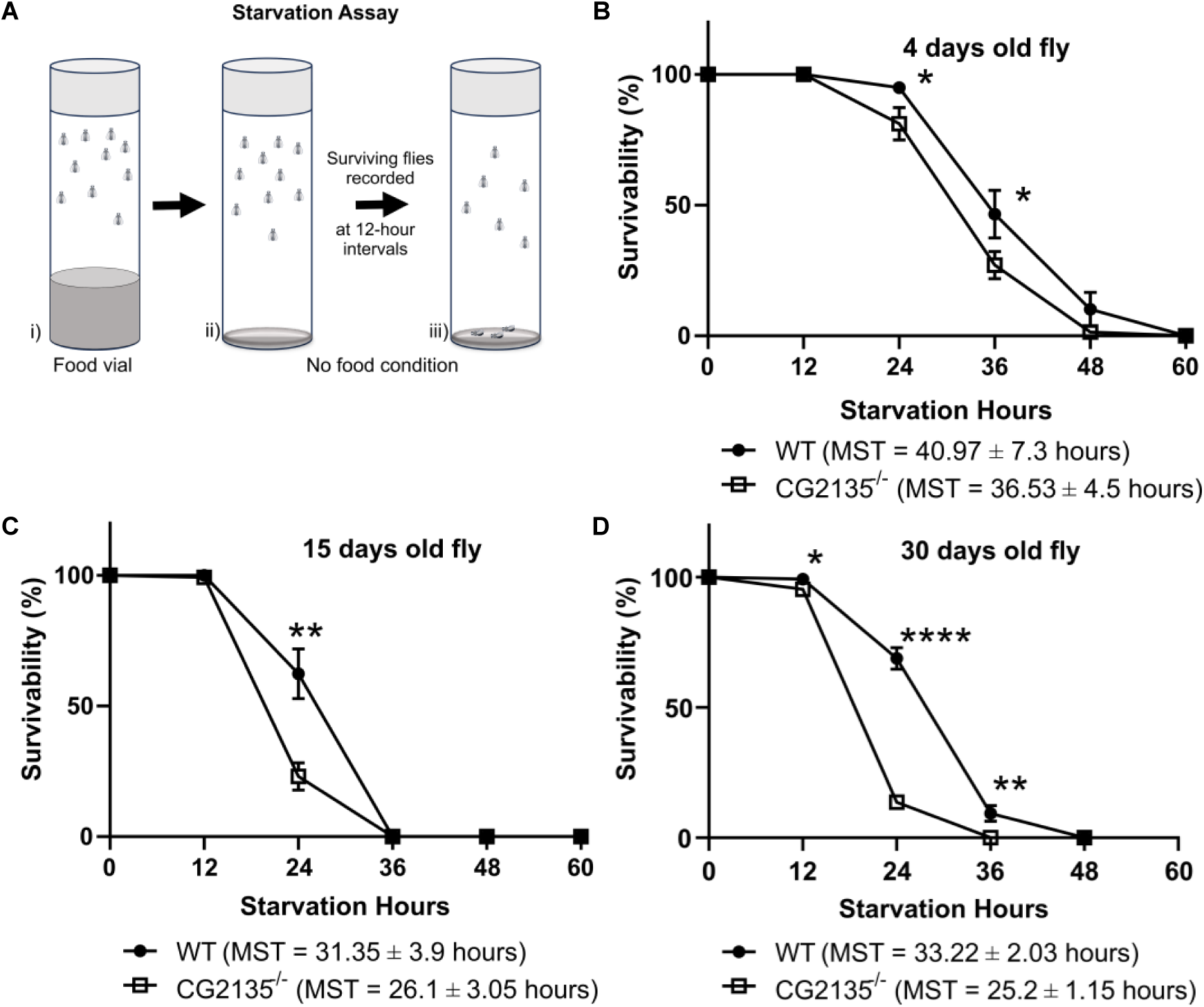
CG2135^-/-^ flies are significantly more susceptible to starvation. (A) Schematic representation of experimental setup for starvation assay. (B-D) Wild type (WT) and CG2135^-/-^ flies of different ages, as indicated in the figure, were subjected to starvation for varying duration and survival curves were plotted. The filled circles and empty squares represent WT the CG2135^-/-^, respectively. Mean survivability time (MST) are mentioned at the bottom of each curve. The survival curves as well as the MST values indicate that the CG2135^-/-^ flies were significantly more susceptible to starvation than their wild type counterparts. All experiments were done independently (N=3), total flies analysed: n≥130. Error bars represent the standard error of mean (SEM). MST means mean survivability time (hours). Asterisks represent level of significance (\*\*\*\**P* ≤ 0.0001, \*\**P* ≤ 0.01, \**P* ≤ 0.05; Student’s *t*-test).

**Figure 2:**
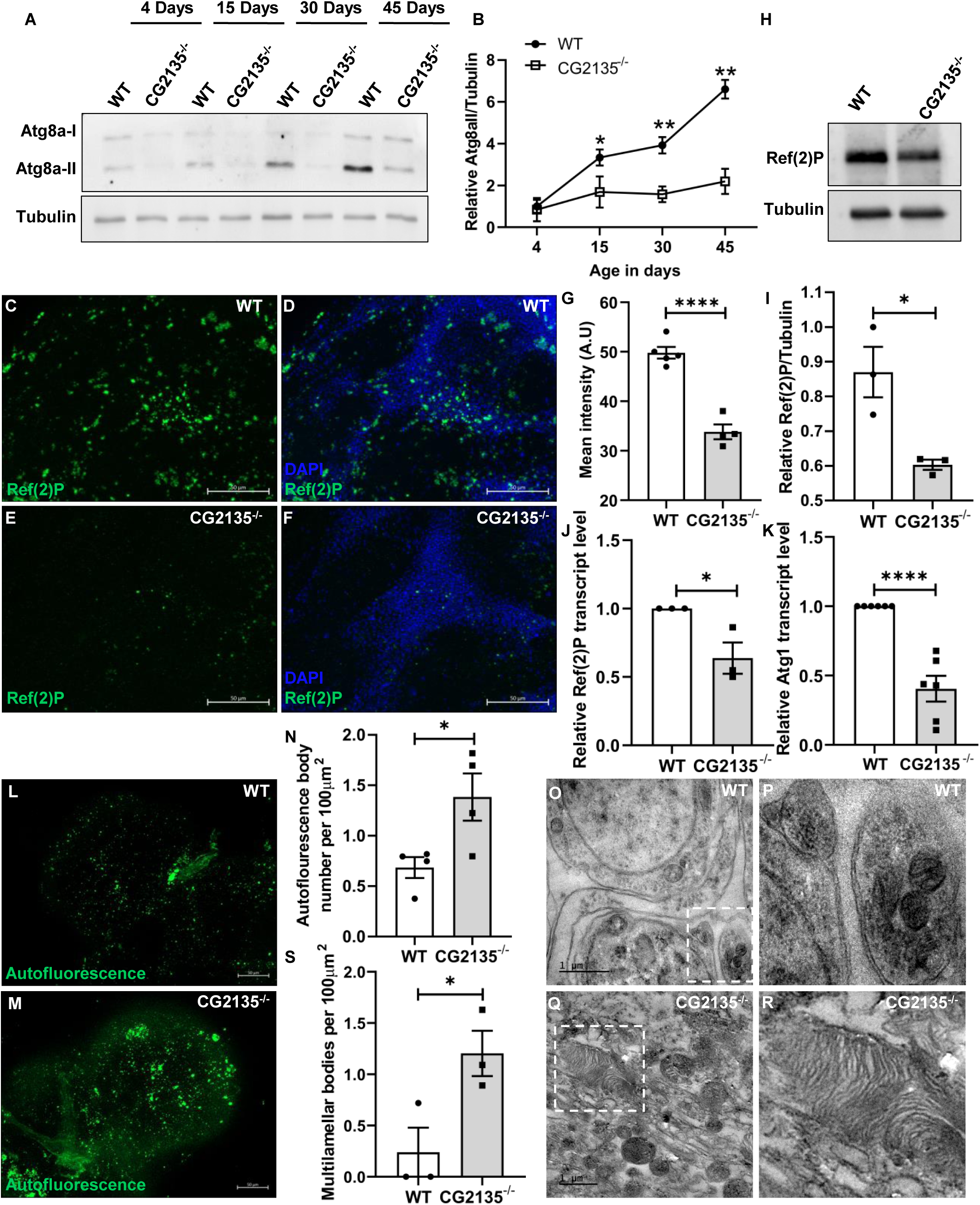
Defective autophagy-lysosome clearance machinery in the CG2135^-/-^ fly brain. (A) Western blot of Atg8a showing the protein level of Atg8a-II (14 kDa) from head lysate of wild type (WT) and CG2135^-/-^ of different ages, as indicated in the figure. α-Tubulin (54 kDa) was used as loading control. (B) The graph represents quantification for Atg8a-II/Tubulin band intensity showing reduced level of Atg8a-II in the CG2135^-/-^ head lysate. (N=3) (C-F) Immunostaining of Ref(2)P in the 30 days old mid-brain of WT and CG2135^-/-^ with anti- Ref(2)P (green) showing reduced Ref(2)P aggregates in the CG2135^-/-^ brain. Nucleus was counterstained with DAPI (blue). (G) The bar graph depicting mean fluorescence intensity of Ref(2)P of WT and CG2135^-/-^ brain (N≥4 fly). (H) Western blot of Ref(2)P (100 kDa) using anti-Ref(2)P from 30 days old WT and CG2135^-/-^ head lysate showing reduced level of the protein in CG2135^-/-^. α-Tubulin (54 kDa) was use as loading control. (I) The bar graph showing the relative level of Ref(2)P/Tubulin protein (N=3). (J) Ref(2)P transcript level in 30 days old WT and CG2135^-/-^ fly head was determined by RT-qPCR and normalised to RP49 as housekeeping gene control. (N=3) (K) Atg1 transcript levels in 30 days old WT and CG2135^-/-^ fly head was determined by RT-qPCR and normalised to RP49 as housekeeping gene control. (N=6) (L-M) Autofluorescent bodies (detected using the FITC channel) in 45 days old WT and CG2135^-/-^ brain showing an increased autofluorescence in CG2135^-/-^. (N) Bar graph depicting mean autofluorescence intensity of WT and CG2135^-/-^ brain (N=4). (O-R) Electron micrograph showing autophagosome and multilamellar bodies from 45 days old WT and CG2135^-/-^ brain. P and R represents magnified images from the respective insets. (S) Quantification of the number of multilamellar bodies compared between WT and CG2135^-/-^ brain has been represented as a bar graph. Multiple fields were analysed from 3 brains of each genotype. The CG2135^-/-^ had increased number of multilamellar bodies. All experiments were done independently. Error bars represent the standard error of mean (SEM). Asterisks represent a level of significance (\*\*\*\**P* ≤ 0.0001, \*\**P* ≤ 0.01, \**P* ≤ 0.05; Student’s *t*-test).

### Abnormal accumulation of enlarged and damaged mitochondria in the CG2135^-/-^ fly brain

One of the unexplained pathological features of the CG2135^-/-^ fly brain is the increased accumulation of MitoTracker-stained puncta ^10^. Upon further analysis, we noted that in the 30-days old CG2135^-/-^ fly brain there was a significant increase in the MitoTracker staining intensity (by ∼ 1.9 folds) as well as the size of the MitoTracker-positive puncta (by ∼ 1.3 folds) compared to the wild type fly brain (Figures 3A-3F). The level of the mitochondrial protein ATP5A was also significantly elevated in the CG2135^-/-^ fly brain, as revealed from our western blot analysis (Figure S2). Taken together, these data indicated an increased abundance of mitochondrial mass in the CG2135^-/-^ fly brain. To gain deeper insights, we next performed ultrastructure analysis of the mitochondria in the 45-days old fly brains using TEM (Figures 3G and 3H). Examination of the TEM images revealed that compared to the wild type fly brain, there was ∼ 1.6 folds increase in the average mitochondria size in the CG2135^-/-^ fly brain (0.27 μm^2^ vs. 0.43 μm^2^) (Figure 3I). Percent frequency distribution analysis of the mitochondria size further revealed that the smaller mitochondria (< 0.2 μm^2^) were much more abundant in the wild type fly brain compared to their CG2135^-/-^ counterparts (∼ 46% in the wild type vs. ∼ 17% in the CG2135^-/-^ fly brain). In contrast, the CG2135^-/-^ fly brain had an increased number of enlarged mitochondria of sizes ≥ 0.4 μm^2^ (∼ 21% in the wild type vs. ∼ 44% in the CG2135^-/-^ fly brain). Abundance of intermediate sized mitochondria (0.2 μm^2^ - 0.4 μm^2^) was also slightly higher in the CG2135^-/-^ fly brain than the wild type (Figure 3J). Aged mitochondria often exhibit distorted cristae structure and are found to be functionally compromised ^38^. This prompted us to methodically examine the mitochondrial ultrastructure from the TEM images and categorized them into healthy, intermediate-damaged, and damaged mitochondria based on the integrity of their cristae structure (Figures 3K-3M). Our analysis revealed that in the wild type fly brain, over 70% of the mitochondria were healthy with orderly packed cristae, whereas, the percentage of healthy mitochondria in the CG2135^-/-^ fly brain was only ∼ 45%. In contrast, the CG2135^-/-^ fly brain exhibited a higher abundance of the damaged and intermediate-damaged mitochondria compared to their wild type partners (damaged: 30% in the CG2135^-/-^ vs. 14% in the wild type, intermediate damaged: 25% in the CG2135^-/-^ vs. 14% in the wild type) (Figure 3N). Having confirmed the abundance of enlarged and damaged mitochondria in the CG2135^-/-^ fly brain, our next goal was to understand its cause and functional consequences.

**Figure 3:**
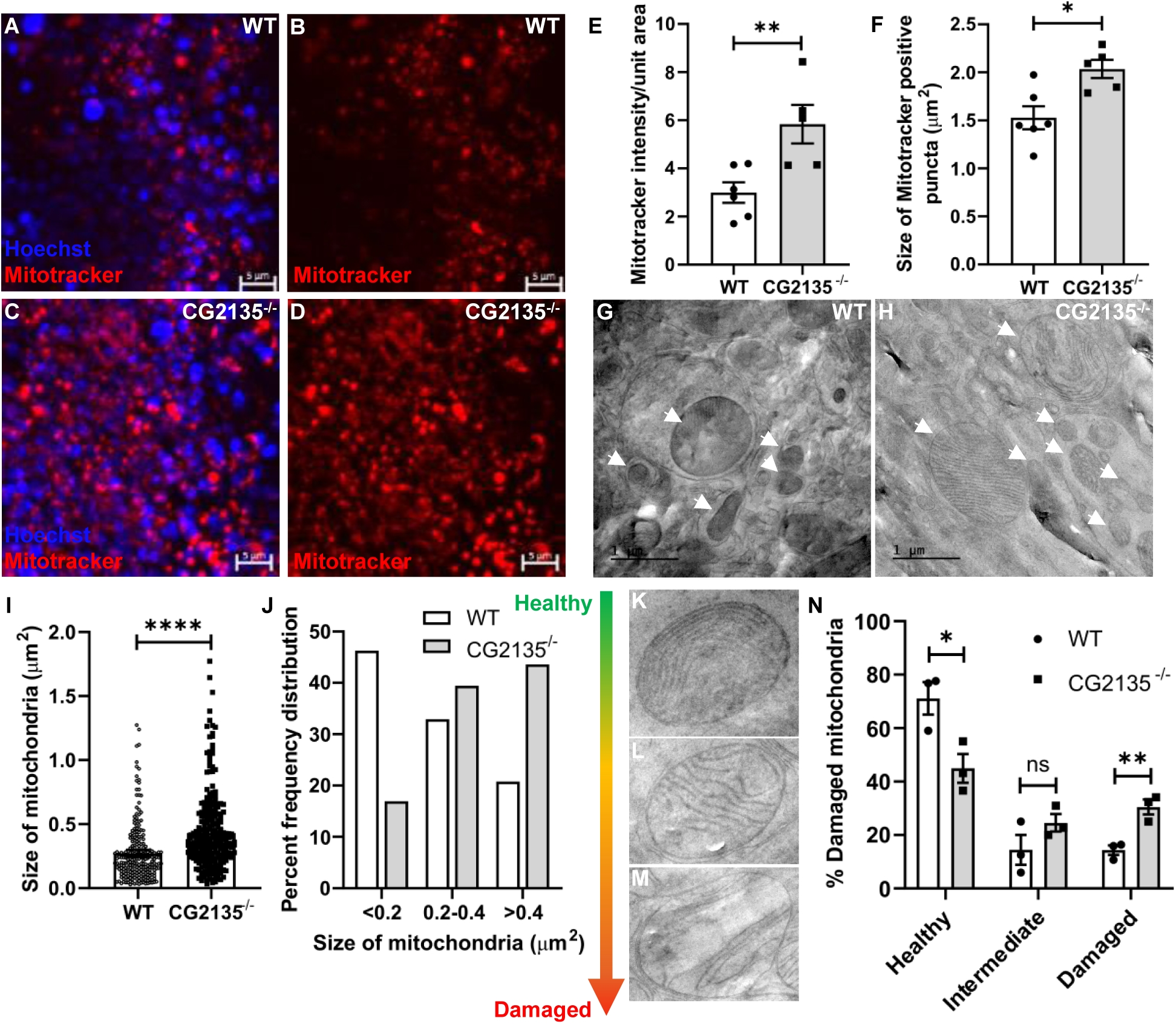
Accumulation of enlarged and damaged mitochondria in the CG2135^-/-^ fly brain. (A-D) Optic lobes of 30 days old wild type (WT) and CG2135^-/-^ fly brain stained with MitoTracker red (showing mitochondria in red) and Hoechst (marking the nucleus in blue), indicating mitochondrial accumulation CG2135^-/-^ fly brain. (E-F) Quantification of MitoTracker intensity and the size of the MitoTracker positive puncta showing increased mitochondrial mass and puncta size in CG2135^-/-^ fly brain (N ≥ 5 fly brain). (G-H) Electron micrographs of 45 days old WT and CG2135^-/-^ fly brains. The mitochondria are marked with white arrows are showing the increased number of mitochondria can be seen in CG2135^-/-^ brain. (I-J) Quantification of the individual mitochondrial area and percent frequency distribution of different sizes showing abundance of larger mitochondria in the CG2135^-/-^ fly brain (N=276 mitochondria from 3 independent experiments) compared to age-matched WT fly brain (N=245 mitochondria from 3 independent experiments). Multiple fields were analysed for measuring the size of the mitochondria using ImageJ. (K-M) Mitochondria observed under TEM were categorized as healthy (K), intermediate (L), and completely damaged (M) based on their cristae structure. (N) The percentage of healthy, intermediate-damaged, and completely damaged mitochondria was quantified from multiple fields of 3 independent experiments, showing reduced healthy but increased damaged mitochondria in the CG2135^-/-^ brain. All experiments were done independently. Error bars represent the standard error of mean (SEM). Asterisks represent a level of significance (\*\*\*\**P* ≤ 0.0001, \*\**P* ≤ 0.01, \**P* ≤ 0.05; Student’s *t*- test).

### Impaired mitophagy in the CG2135^-/-^ fly brain

Mitochondria are particularly susceptible to oxidative damage due to continuous ROS production during cellular respiration. As a cellular cleansing mechanism, the damaged mitochondria are eliminated through the process of mitophagy, a specialized form of autophagy ^17,18^. In this context, it is important to note that impairment of mitophagy has been reported in several age-related neurodegenerative diseases ^19^. These prior reports, combined with our finding that autophagy is suppressed in the CG2135^-/-^ fly brain, led us to investigate whether these flies also suffer from a mitophagy defect. For this, 45 days old fly brains were co-stained with LysoTracker (green) and MitoTracker (red), and the mitophagy status was analysed by quantifying the extent of colocalization of lysosomes and mitochondria from confocal images. Data presented in Figures 4A-4H clearly shows that, compared to the wild type flies, there was a significantly reduction in the lysosome-mitochondria colocalization in the CG2135^-/-^ fly brain, indicating a mitophagy defect in these flies. This was further corroborated by our TEM images, which showed fewer autophagosome-engulfed mitochondria in the 45-day-old CG2135^-/-^ fly brain compared to their wild type counterparts (Figures 4J and 4L). As an alternative approach to analyse mitophagy, we isolated the mitochondria-rich fraction from 45 days old fly brains and checked for the presence of Atg8a -II, an autophagosome-associated protein, in that fraction. Our western blot data showed that the Atg8a -II protein levels in the mitochondria-rich fraction isolated from the CG2135^-/-^ fly brain was significantly lower than in the age-matched wild type flies (Figures 4M-4O). Taken together, our data suggest that impaired mitophagy is the reason behind the accumulation of enlarged and damaged mitochondria in the CG2135^-/-^ fly brain. This prompted us to investigate the mitochondrial function in the CG2135^-/-^ fly brain, particularly in terms of ATP production.

**Figure 4:**
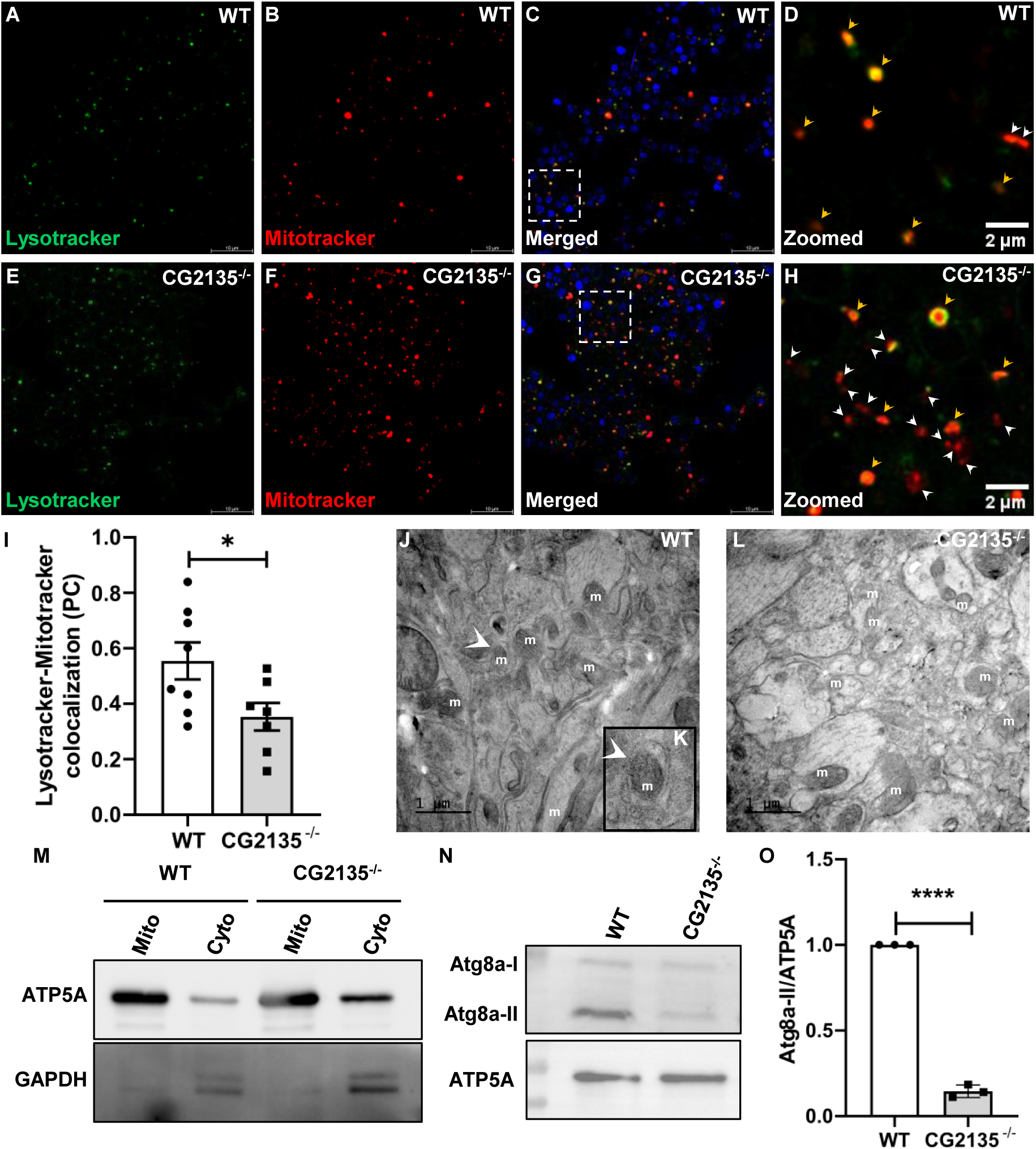
Impaired mitophagy in the CG2135^-/-^ fly brain. (A-H) MitoTracker-LysoTracker co-staining of 45 days old wild type (WT) and CG2135^-/-^ fly brain (optic lobe region) showing fewer colocalization spots (yellow puncta, indicated with yellow arrows) in the CG2135^-/-^ fly, indicating reduced mitophagy. Only MitoTracker positive red puncta are indicated with white arrows. Hoechst is (blue) used for counterstaining the nucleus. (I) Pearsons’ colocalization coefficient (PCC) analysis showing reduced LysoTracker-MitoTracker colocalization in the CG2135^-/-^ brain compared to WT (N ≥ 7). (J-L) Electron micrograph of 45 days old WT and CG2135^-/-^ fly brains showing reduced mitochondria-sequestered autophagy vesicle (K) in the CG2135^-/-^ fly brain as indicated by white arrow. (M) Verification of mitochondria-rich fraction isolated from the 45 days old fly head by western blot for mitochondrial protein, ATP5A (55 kDa), and cytosolic marker GAPDH (36 kDa). (N) The western blot with anti-Atg8a showing reduced Atg8a-II protein in the mitochondria-rich fraction from the CG2135^-/-^ fly head. ATP5A was used as the loading control. (O) The bar graph showing the quantification for the Atg8a-II/ATP5A. N=3 All experiments were done independently. Error bars represent the standard error of mean (SEM). Asterisks represent a level of significance (\*\*\*\**P* ≤ 0.0001, \**P* ≤ 0.05; Student’s *t*-test).

### CG2135^-/-^ fly suffers from brain energy deficit

Mitophagy plays a neuroprotective role by maintaining mitochondrial homeostasis, thereby ensuring sufficient ATP supply for proper brain function ^39,40^. Since mitochondrial membrane potential (ΔΨm) is a key indicator of functionally active mitochondria and is essential in driving ATP synthesis, we used JC-1 dye to analyse ΔΨm in the mitophagy-deficient 45 days old CG2135^-/-^ fly brain. JC-1 is a lipophilic green fluorescent monomeric dye that accumulates within the mitochondria in a potential-dependant manner and forms aggregates (J-aggregate) shifting its fluorescence property to red. The red-to-green fluorescence ratio of the JC-1 dye is often used to measure to ΔΨm ^41^. Compared to the age-matched wild type flies, we observed > 40% reduction in the red-to-green fluorescence ratio in the CG2135^-/-^ fly brain, indicating a significant decrease in ΔΨm in these flies (Figures 5A-5G). Strikingly, the ATP levels in the 45 days old CG2135^-/-^ fly brain was found to be reduced by >50% than in the wild type fly brain (Figure 5H). This data indicates that the older CG2135^-/-^ flies suffer from severe brain energy deficit, which may be the trigger for neurodegeneration in these flies. ATP is produced in the mitochondrial inner membrane by the enzyme ATP synthase, which utilizes the proton gradient across the mitochondrial membrane as the driving force for generating ATP ^41^. It has been reported that if ΔΨm is compromised for some reason, ATP synthase reverses its direction acting as an ATP-consuming enzyme to translocate protons from the mitochondrial matrix to the intermembrane space. In such situations, the expression of ATPase Inhibitory Factor 1 (ATPIF1) is upregulated to prevent wasteful ATP expenditure by inhibiting the ATP synthase activity ^42^. Consistent with these findings, we observed a significant upregulation of ATPIF1 transcript level in the 45 days old CG2135^-/-^ fly brain, suggesting that ATP synthase activity is indeed inhibited in these flies, possibly to counteract further depletion of ATP levels (Figure 5I).

**Figure 5:**
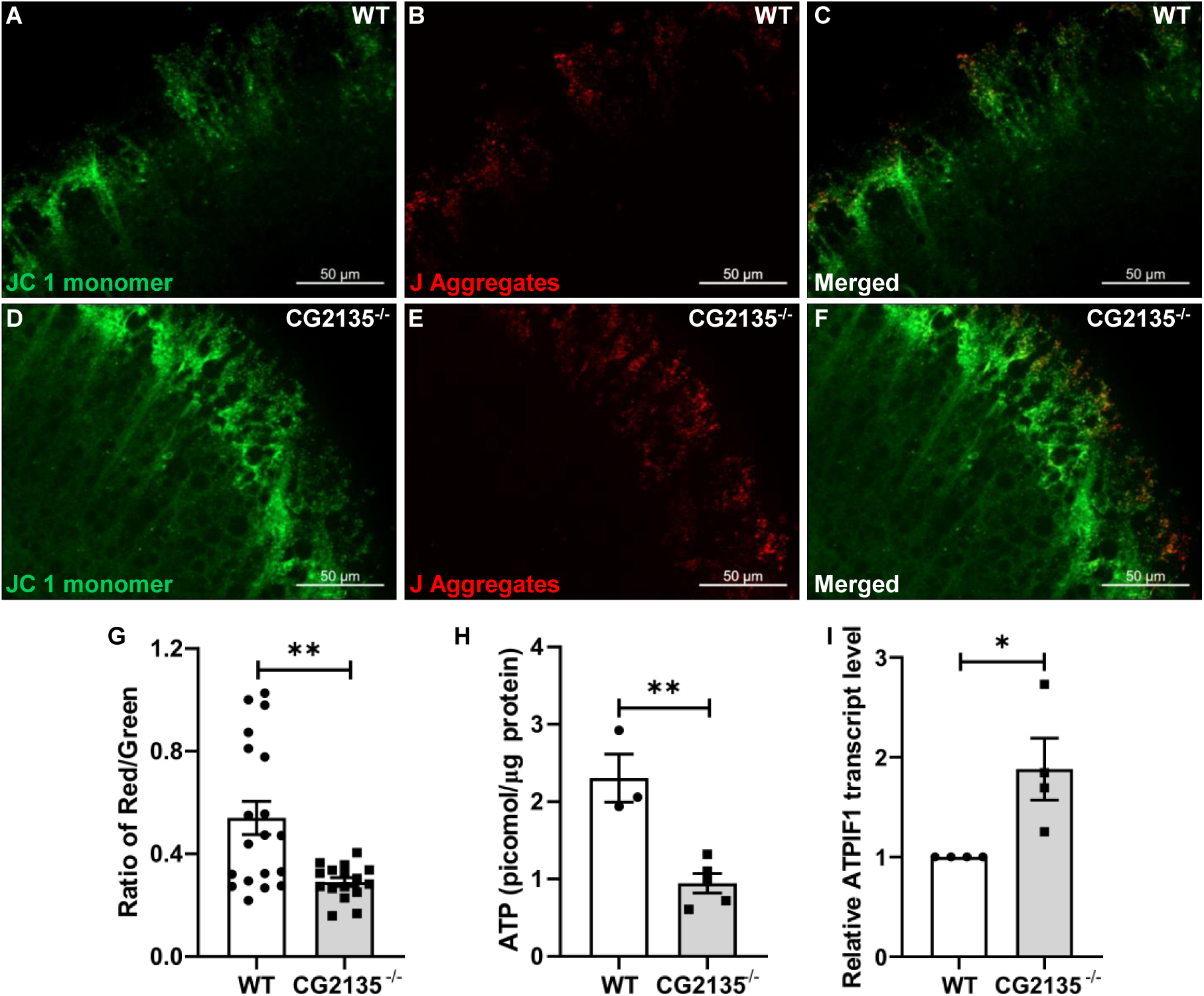
Reduced mitochondrial membrane potential and ATP levels in the CG2135^-/-^ fly brain. (A-F) The optic lobes of 45 days wild type (WT) and CG2135^-/-^ fly brains were stained with JC-1 dye for assessing the mitochondrial membrane potential (ΔΨm). (G) The bar graph represents quantification of the red/green ratio showing reduced ΔΨm in CG2135^-/-^ brain (N ≥ 16) (H) The bar graph represents quantification of total ATP from 45-days old fly head (N = 3), showing significantly reduced ATP levels in the CG2135^-/-^ fly brains. (I) Relative transcript levels of ATPIF1 in 45 days old CG2135^-/-^ fly head determined by RT-qPCR and normalised to RP49 as housekeeping gene control (N=4). All experiments were done independently. Error bars represent the standard error of mean (SEM). Asterisks represent a level of significance (\*\**P* ≤ 0.01, \**P* ≤ 0.05; Student’s *t*-test).

### Increased apoptosis in the CG2135^-/-^ fly brain

We previously reported prominent signs of neurodegeneration, including loss of dopaminergic neurons, in the CG22135^-/-^ fly brains ^10^. However, the mechanism underlying this neurodegeneration is far from clear. Since mitochondrial damage and ATP depletion have been reported to induce apoptotic cell death in both neuronal and non-neuronal cells ^43–45^, we checked the status of apoptosis in the energy-deficient 45 days old CG2135^-/-^ fly brain. Our western blot data clearly showed that the CG2135^-/-^ fly brains had significantly elevated levels of cleaved-caspase-3, a key mediator of apoptosis (Figures 6A an 6B) ^46^. Interestingly, co-immunostaining of the fly brains with antibodies against cleaved-caspase-3 and ELAV (a pan-neuronal marker) ^47^ revealed that both neuronal (ELAV-positive) as well as non-neuronal (ELAV-negative) cells in the CG2135^-/-^ fly brain showed signs of apoptosis (Figure S3). These findings were corroborated by transcript analysis of the proapoptotic factor Hid^48^, which showed ∼ 2.4 folds upregulation in the CG2135^-/-^ fly brain compared to their age-matched counterparts (Figure 6C). For direct visualization of DNA fragmentation in apoptotic cells, we next performed TUNEL staining in whole mount brains. We observed ∼ 2.8 folds increase in the abundance of TUNEL-positive cells in the 45 days old CG2135^-/-^ fly brains compared to the wild type fly brains (Figures 6D-6L). When the brains were co-stained with anti-Tyrosine Hydroxylase antibody (to mark the dopaminergic neurons) and TUNEL (to mark the apoptotic nuclei), we observed that ∼ 18% of the dopaminergic neurons in the CG2135^-/-^ fly brains were TUNEL positive, as compared to only ∼ 2% in the wild type flies (Figures 6M-6S). Collectively, these data suggest that apoptosis plays a key role in neurodegeneration in the MPS VII fly.

**Figure 6:**
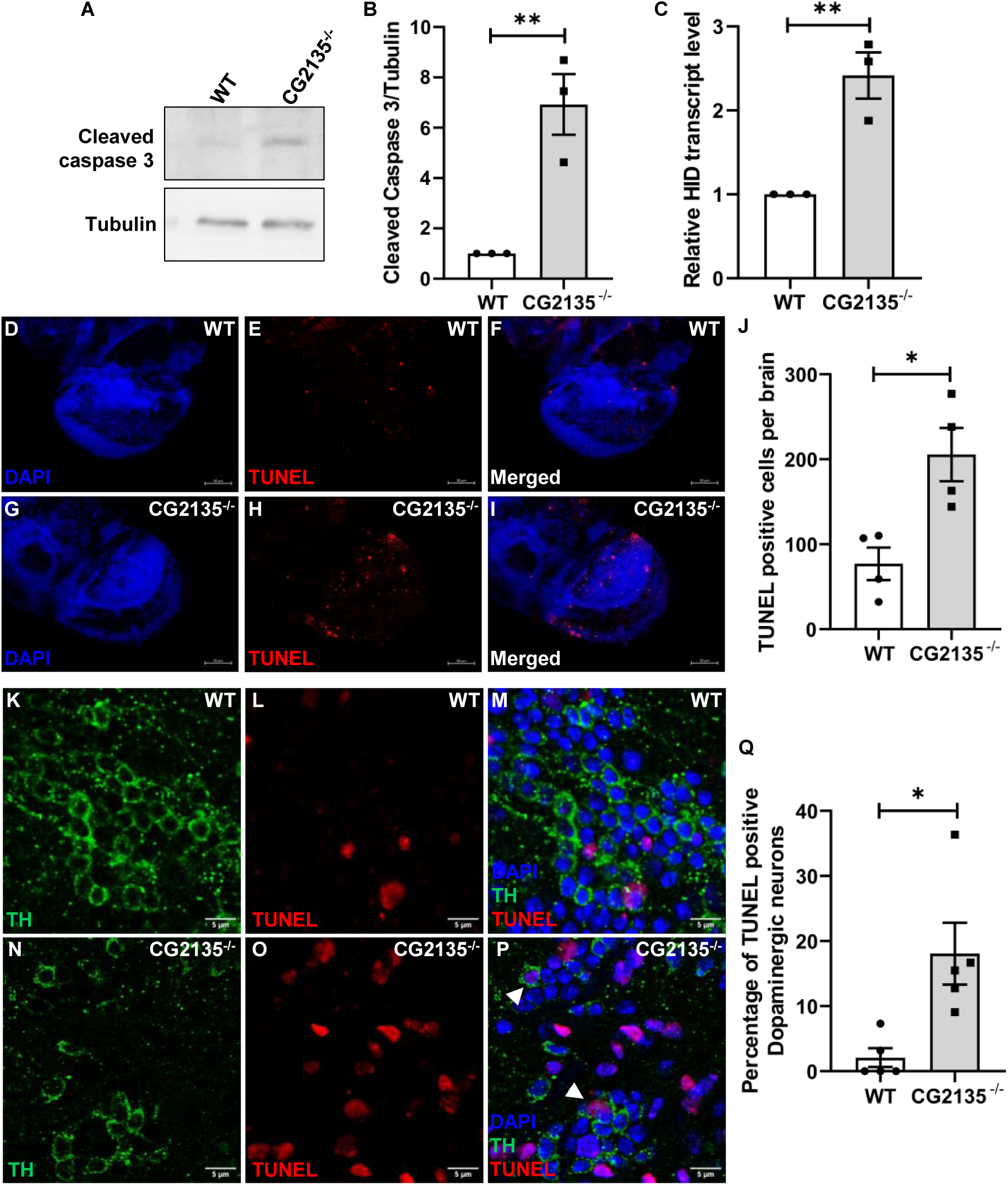
Increased apoptosis in the CG2135^-/-^ fly brain. (A) Western blot of Cleaved-caspase 3 using anti-Cleaved-caspase 3 antibody showing increased processed form of 22 kDa at 45 days old CG2135^-/-^ fly head compared to the wild type (WT). α- Tubulin (54 kDa) was used as the loading control. (B) The bar graph showing the relative level of Cleaved-caspase 3/Tubulin. N=3 (C) Relative transcript level of pro-apoptotic gene Hid determined by RT-qPCR and normalised to RP49 as housekeeping gene control (N=3) (D) TUNEL staining of 45 days old WT and CG2135^-/-^ brain, DAPI (blue) used for marking the nucleus. (J) The bar graph showing quantification of the TUNEL-positive nucleus showing increased TUNEL-positive cells in CG2135^-/-^ fly brain. (N=4 fly brain) (K-P) Co-staining of TUNEL and anti-TH (tyrosine hydroxylase) showing elevated TUNEL-positive dopaminergic neurons in 45 days old CG2135^-/-^ brain. DAPI (blue) was used for counter- staining the nucleus. (Q) The bar graph representing quantification of apoptotic dopaminergic neurons (TUNEL and TH double positive) in WT and CG2135^-/-^. (N=5) All experiments were done independently. Error bars represent the standard error of mean (SEM). Asterisks represent level of significance (\*\**P* ≤ 0.01, \**P* ≤ 0.05; Student’s *t*-test).

### Resveratrol treatment corrects mitophagy defect and prevents ATP loss in the CG2135^-/-^ fly brain

We previously reported that resveratrol treatment reversed the locomotor deficit in the CG2135^-/-^ fly and protected against loss of their dopaminergic neurons ^10^. But how these abnormalities were corrected by resveratrol remained a mystery. To evaluate the mechanism of resveratrol action, we treated the CG2135^-/-^ flies with it for 30-45 days starting from day 4 post-eclosion. We then assessed the status of Atg1 transcript and the level of mitochondrial protein ATP5A in the brain. We found that Atg1 transcript level, which was significantly downregulated in the CG2135^-/-^ fly brain, was completely restored to the normalcy in the resveratrol-treated CG2135^-/-^ flies (Figure 7A). The elevated ATP5A level in the CG2135^-/-^ fly brain was also reduced to the to the basal level upon resveratrol treatment (Figures 7B and 7C). Most importantly, resveratrol treatment protected the CG2135^-/-^ flies from brain energy deficit by completely restoring their ATP levels (Figure 7D). From our data, it can be conjectured that correction of mitophagy defect and restoration of ATP levels in the CG2135^-/-^ fly brain upon resveratrol treatment is the mechanistic basis for its neuroprotective effect.

**Figure 7:**
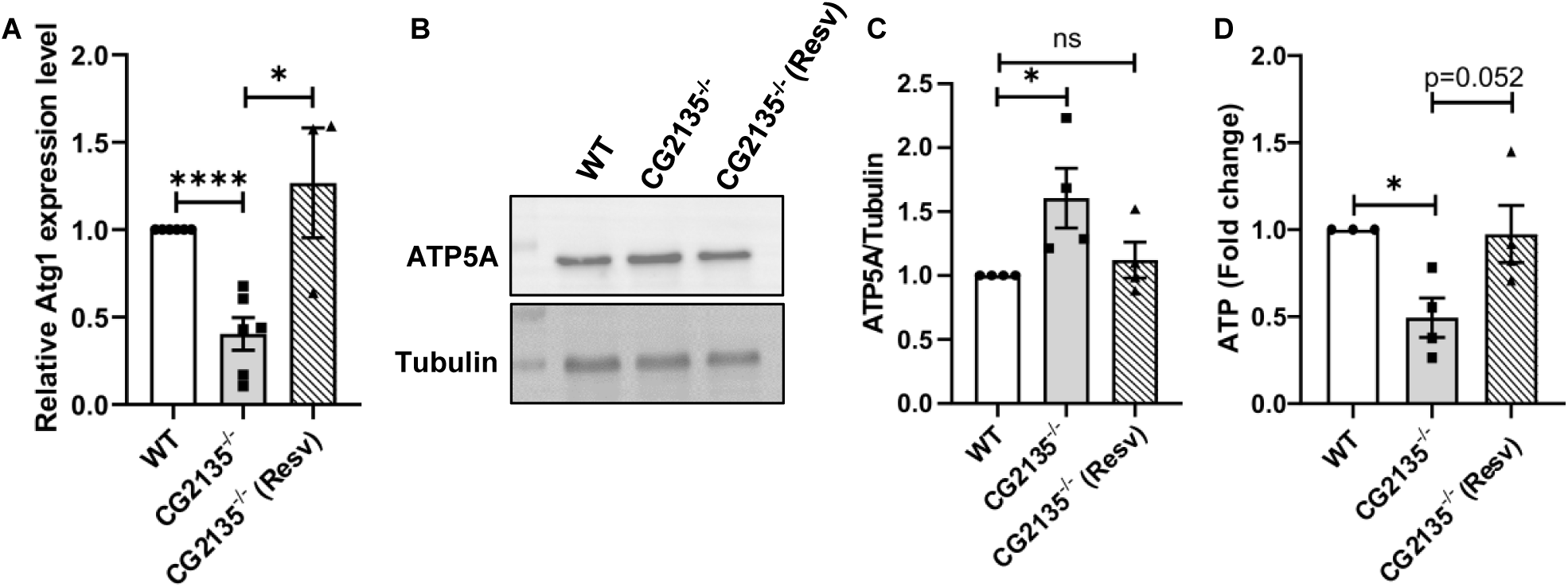
Resveratrol treatment corrects mitophagy defect and prevents ATP loss in the CG2135^-/-^ fly brain. (A) The bar graph represents the Atg1 transcript level determined by RT-qPCR in 30 days old wild type (WT), CG2135^-/-^ and resveratrol-treated CG2135^-/-^ fly head showing restoration of Atg1 level on resveratrol treatment. The gene expression was normalised to RP49 as housekeeping gene control. (N=6 for untreated WT and CG2135^-/-^; N=3 for resveratrol treated CG2135^-/-^). (B) Western blot of ATP5A using anti-ATP5A antibody showing restoration of the protein level in the head lysate of CG2135^-/-^ fly compared to untreated WT and CG2135^-/-^ at 45 days. (C) The bar graph showing quantification of ATP5A/Tubulin protein level in untreated WT, CG2135^-/-^ and resveratrol-treated CG2135^-/-^ fly head. (N=4). (D) The bar graph represents the ATP level of WT, CG2135^-/-^ and resveratrol-treated CG2135^-/-^ fly head at 45 days. ATP level in CG2135^-/-^ fly head restored to normalcy with resveratrol treatment. (N ≥ 3) All experiments were done independently. Error bars represent the standard error of mean (SEM). Asterisks represent a level of significance (\*\*\*\**P* ≤ 0.0001, \*\*\**P* ≤ 0.001, \**P* ≤ 0.05; Student’s *t*- test).

## Discussion

Despite being known as a debilitating disease with prominent neurological symptoms, the mechanism of neurodegeneration in MPS VII remains poorly understood. The work presented here aimed to bridge this knowledge gap by employing a *Drosophila* model of the disease (the CG2135^-/-^ fly). We report a gross defect in the autophagy-lysosome clearance pathway in the CG2135^-/-^ fly brain, which also manifested as impaired mitophagy. As a consequence, there was an accumulation of structurally deformed and functionally compromised mitochondria leading to bioenergy deficit in the brain and apoptotic neurodegeneration. We also demonstrated that mitophagy defect and brain energy deficit in the CG2135^-/-^ fly could be corrected by treatment with resveratrol, providing a potential explanation for its neuroprotective property.

Autophagy is crucial for maintaining cellular homeostasis by clearing cytotoxic protein aggregates and damaged organelles, a process especially vital for post-mitotic cells like neurons in the central nervous system (CNS) ^22,49^. In fact, defective autophagy has been linked with neurodegenerative diseases like Alzheimer’s, Parkinson’s, and ALS ^49^. Lysosomal storage disorders (LSDs) such as multiple sulfatase deficiency, Gaucher disease and mucolipidosis type IV, often exhibit neurodegeneration in brain with impaired autophagy ^14,15,34^. Despite these studies, the status of autophagy in the MPS VII brain remained completely unexplored. Even for other MPS disorders, there is hardly any systematic study on brain autophagy. We, therefore, took up this challenge following the lead from our earlier observation of defective autophagy in the CG2135^-/-^ larvae ^20^. Also, in the present study, the adult CG2135^-/-^ flies were found to be highly susceptible to starvation. Since autophagy is typically induced as a survival response under nutrient starvation, this starvation-sensitivity was an indicator of a global autophagy defect in the CG2135^-/-^ flies ^26,27^. However, as autophagic response is highly tissue-specific and it is known that starvation-induced autophagy in brain is much less pronounced than in other tissues ^50^, it was imperative for us to specifically analyze the autophagy status in the adult CG2135^-/-^ fly brains. A significantly reduced abundance of Atg8a-II protein, a bona fide marker of autophagosomes, confirmed that autophagy is indeed altered in the CG2135^-/-^ fly brains. It was interesting to note that the Atg8a-II levels in the wild type fly brains gradually increased with age, indicating normal autophagy induction but progressive decline of lysosome-mediated autophagosome turnover leading to autophagosome accumulation ^51,52^. But, the Atg8a-II levels in the CG2135^-/-^ fly brains did not increase with age and remained almost at the basal level throughout. This result, together with our data demonstrating significantly reduced expression of Atg1 and Ref(2)P in the CG2135^-/-^ fly brains, suggest impaired autophagy induction in the MPS VII flies. The process of autophagy is fine-tuned by multilayered regulation of Atg1/ULK1 initiation factor ^53^. Various transcription factors (like ATF4, FoxO3, STAT1, and ZKSCAN3) and signal transducers (like mTOR and AMPK) have been shown to regulate the expression and activity of Atg1/ULK1^54,55^. It remains to be seen whether altered expression of any of these regulatory proteins is responsible for downregulation of Atg1 in the CG2135^-/-^ fly brains. Atg1/ULK1, once activated, initiates autophagy by phosphorylating its downstream targets leading to the assembly of Atg12-Atg5-Atg16L1 complex. It acts as a E3 ligase, facilitating lipidation of cytosolic Atg8a-I/LC3-1 to autophagosome-bound Atg8a-II/LC3-II ^51^. So, it is not surprising that under the condition of suppressed Atg1 expression, Atg8a-II levels remained consistently low at all ages in the CG2135^-/-^ fly brains. The low transcript as well as protein levels of Ref(2)P in the CG2135^-/-^ fly brains was rather intriguing. Ref(2)P, the fly orthologue of the mammalian autophagic cargo adaptor protein p62, usually accumulates when there is a block in the autophagy pathway ^56^. However, a contrasting observation was reported in peripheral blood mononuclear cells (PBMCs) from Gaucher disease patients, where LC3-II levels as well as p62 transcript were both found to be reduced compared to PBMCs from healthy donor ^57^. This suggest that Ref(2)P/p62 expression and autophagy induction could be simultaneously impaired under certain conditions, although physiological implication of such phenomenon is not clear. Ref(2)P/p62 being a TFEB-inducible gene, its reduced expression in the CG2135^-/-^ fly brains might be a collateral consequence of TFEB downregulation ^58^. While TFEB has been found to be dysregulated in some neurodegenerative diseases ^59^, its status is not known in any of the MPS disorders and this remains a future area for investigation.

In addition to stunted autophagy induction, defective clearance of the autophagosomal content was also evident in the CG2135^-/-^ fly brains, as indicated by accumulation of lipofuscin-like inclusions and multilamellar bodies (MLBs). Lipofuscin is an intralysosomal autofluorescent material mainly composed of lipids and misfolded proteins that are resistant to lysosomal degradation. Lipofuscin accumulation can be aggravated when lysosomal degradation process is hampered, particularly in certain lysosomal storage diseases such as Batten disease, Tay-Sachs disease and Sanfilippo syndrome ^33,60,61^. Accumulation of concentric multilamellar structures in the CG2135^-/-^ fly brains is another signature for dysfunctional lysosomal activity, as has been observed in the brains of a *Drosophila* model of mucolipidosis type IV and fibroblasts form Parkinson’s disease patients ^34,36,62^.

Autophagy being a critical homeostatic machinery in neuronal as well as some non-neuronal cells of the central nervous system ^63–65^, blockage in multiple steps of this pathway in the CG2135^-/-^ fly brains was expected to have deleterious consequences. This notion was validated when we found significantly reduced mitochondria-lysosome colocalization and fewer autophagosome-sequestered mitochondria in the CG2135^-/-^ fly brains, indicating a defect in mitophagy, a specialized form of autophagy to remove damaged mitochondria. As the primary site for cellular energy production, mitochondria are prone to oxidative damage ^18^. This vulnerability is even more pronounced in high energy-demanding tissues like the brain ^66^. To maintain a healthy pool of mitochondria, cells are endowed with a robust mitochondrial quality control mechanism that comprises two opposing but coordinated processes: mitochondrial biogenesis, and removal of damaged mitochondria through mitophagy ^17,18^. The canonical mitophagy is mediated by the PINK1-Parkin signalling cascade. In this pathway, PINK1 is stabilized on the mitochondrial outer membrane, following which it phosphorylates and recruits Parkin, thereby triggering ubiquitin-dependent mitophagy ^17^. In PINK1-Parkin-independent non-canonical mitophagy, adaptor molecules on the mitochondrial membrane, such as FUNDC1, can directly initiate mitophagy ^67,68^. It has been reported that Atg1/ULK1 plays a vital role in regulating mitophagy through both canonical and non-canonical pathways by phosphorylating key molecules like Parkin or FUNDC1 ^68–71^. Thus, it can be speculated that reduced Atg1 expression in the CG2135^-/-^ fly brains is responsible, at least in part, for the observed mitophagy defect. Decreased Ref(2)P levels may also contribute to impaired mitophagy, as p62/Ref(2)P downregulation has been shown to cause autophagic cargo loading failure ^72^. Furthermore, PINK1-Parkin-mediated mitophagy is reported to depend on p62/Ref(2)P ^73^, suggesting that reduced levels of this cargo adapter protein could aggravate the mitophagy defect in the CG2135^-/-^ fly brains.

The mitophagy defect in the CG2135^-/-^ fly brains provided an explanation for the accumulation of enlarged mitochondria with deformed cristae structure and reduced ΔΨm. It is well-documented that such excessive buildup of damaged mitochondria is a direct consequence of a mitophagy defect, regardless of the actual cause of the defect ^74,75^. It is noteworthy that while signs of autophagy defect were evident even in the younger CG2135^-/-^ flies, accumulation of damaged and depolarized mitochondria in their brains could be detected only in the older flies, aged 30 days and beyond. In fact, the brains of 45 days old CG2135^-/-^ flies were found to be bioenergetically inefficient showing nearly a 50% drop in ATP levels. Such mitophagy deficiency-induced depletion of ATP levels in the brain has been observed in other neurodegenerative diseases ^76,77^, but not reported earlier in any of the MPS disorders. Since ΔΨm is a key driver for ATP synthesis ^41^, it can be inferred that ATP depletion in the CG2135^-/-^ fly brain is primarily due to accumulation of depolarized mitochondria with significantly reduced ATP-synthesizing ability. Consumption of ATP via reversal of ATP synthase activity, as often occurs in depolarized mitochondria ^42^, may further exhaust the ATP levels in the CG2135^-/-^ fly brains. Robust upregulation of ATPIF1 transcript level in the CG2135^-/-^ fly brains support this possibility. Under conditions of compromised ΔΨm, ATPIF1 is known to act as an inhibitor of ATP synthase activity, preventing ATP wastage ^42^.

Depleted level of intracellular ATP is an established determinant of apoptotic cell death ^45,78^. Thus, it was not unexpected that ATP depletion in a high energy-demanding tissue like brain would induce apoptotic cell death in the CG2135^-/-^ flies. Multiple reports have confirmed that autophagy machinery plays a crucial pro-survival role and defects in this process predispose cells to apoptosis ^79–81^. Therefore, in addition to ATP depletion, autophagy impairment could be another factor for inducing apoptosis in the CG2135^-/-^ fly brains. Earlier, we reported a significant loss of dopaminergic neurons in the CG2135^-/-^ fly brains, which could be mitigated with resveratrol treatment ^10^. However, the underlying cause of this neuronal loss and the mechanism by which resveratrol conferred protection against it remained unclear. The cause of neuronal loss became evident when we discovered that neuronal cells, particularly the dopaminergic neurons, are undergoing apoptosis in the CG2135^-/-^ fly brains. This finding mirrors one of the hallmarks of Parkinson’s disease, which is characterized by progressive loss of dopaminergic neurons via apoptosis ^82^. It is worth noting that dopaminergic neurons have a high metabolic demand due to their extensive axonal network and dopamine secretory activity, making them particularly vulnerable to energy crisis ^83^. Intriguingly, we observed that resveratrol treatment could correct the mitophagy defect and fully restore ATP levels in the CG2135^-/-^ fly brains, providing a mechanistic explanation for its neuroprotective property. These findings align with an earlier report demonstrating that resveratrol provides neuroprotection by safeguarding against rotenone-induced mitochondrial damage through the enhancement of autophagic activity ^84^.

Apart from neuronal cells, signs of apoptosis were also found in some non-neuronal cells of the CG2135^-/-^ fly brains. In *Drosophila* brain, non-neuronal cells are predominantly glial cells, which play crucial roles in maintaining and protecting neural environment ^85^. Whether degeneration of glial cells also drives neurodegeneration in MPS VII remains an open question. It has been observed that degenerating glial cells derived from MPS II mice were able to promote neuronal death in a co-culture condition ^86^. Given that the role of mitophagy in glial cells has been implicated in the pathogenesis of other neurodegenerative diseases ^87,88^, it may be worthwhile to investigate the contributions of these cells in neurodegeneration associated with MPS VII and related disorders.

To conclude, the work presented here significantly advances our understanding of the pathogenesis of MPS VII by uncovering a previously unknown mechanistic link between mitophagy defect, mitochondrial malfunction leading to brain energy deficits, and apoptotic neurodegeneration. This study also led to the identification of critical molecular alterations in the autophagy-lysosomal pathway in the MPS VII fly brains, which may provide novel therapeutic leads for pharmacological interventions. The promising results with resveratrol treatment in correcting the mitophagy defect and preventing loss of ATP could be an important first step towards this direction. Since MPS disorders share many commonalities, our findings are likely to have broader implications in understanding the mechanisms of neurodegeneration associated with this group of lysosomal storage diseases.

## Acknowledgements

This research was supported by the MoE, Govt. of India grant STARS/APR2019/BS/779/FS awarded to RD. The authors express their sincere gratitude to Mohit Prasad and the members of his fly lab for their support in fly maintenance. Thanks are also due to Sankar Maiti, Arnab Gupta, and Jayasri Das Sarma for sharing critical reagents and facilities. The authors acknowledge Sudipta Bar for his valuable suggestions and Subhankar Dolai for his critical comments on the manuscript. Abhrajyoti Nandi, Sujoy Bose and Subhajit Majumdar are appreciated for their technical assistance. The support of the TEM facility at the Institute of Life Sciences (ILS), Bhubaneswar is acknowledged. NM was supported by CSIR fellowship and AD was supported by DBT and ICMR-SRF fellowships.

## Author contributions

N.M. A.D. and R.D. conceptualized the work and designed the experiments; N.M. and A.D. performed the experiments; N.M. A.D. and R.D. analysed the data; N.M. A.D. and R.D. wrote the manuscript; R.D. supervised the work and acquired the funding.

## Supplemental information

## Materials and methods

All the reagents are enlisted in the resource table.

### Key resource table

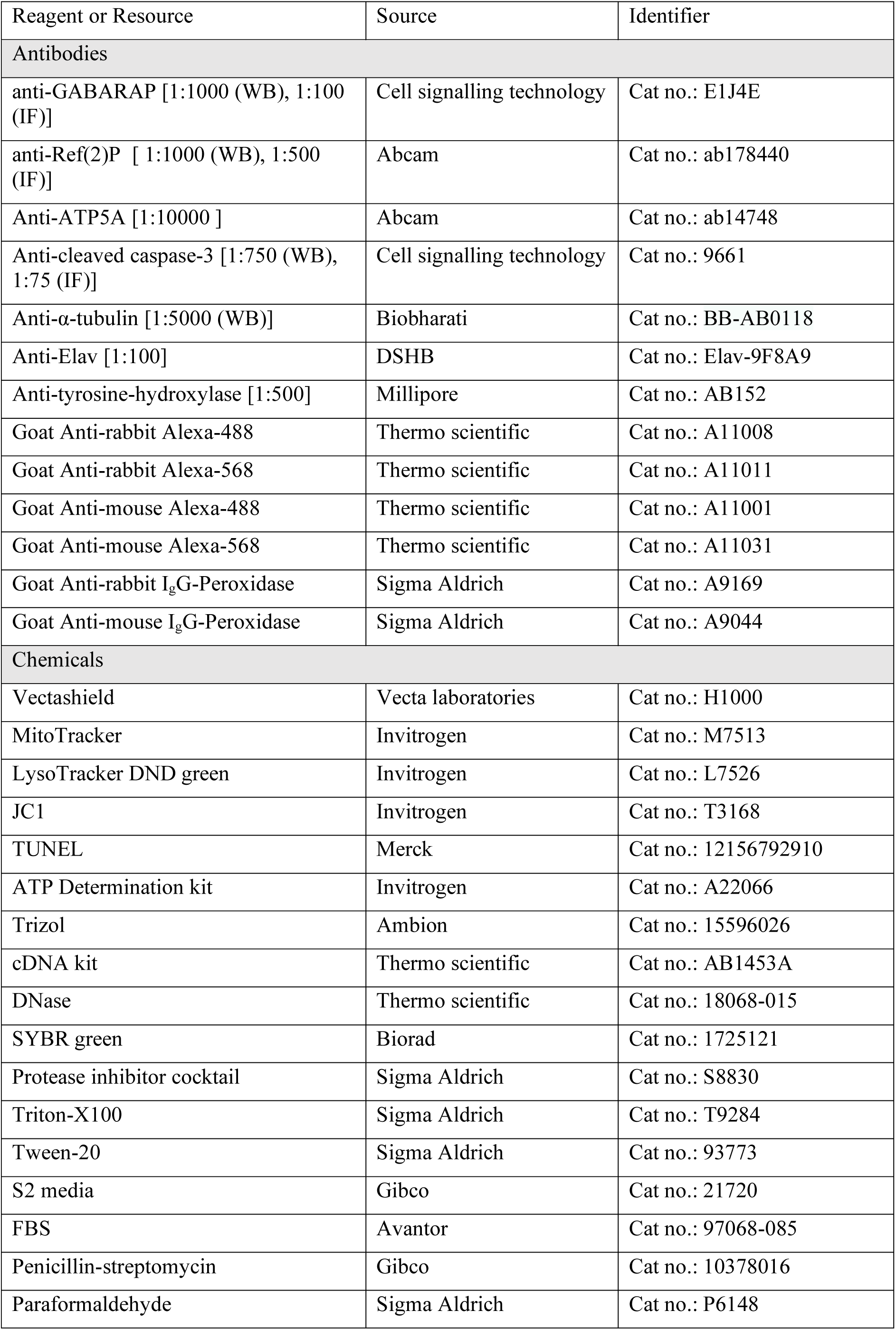
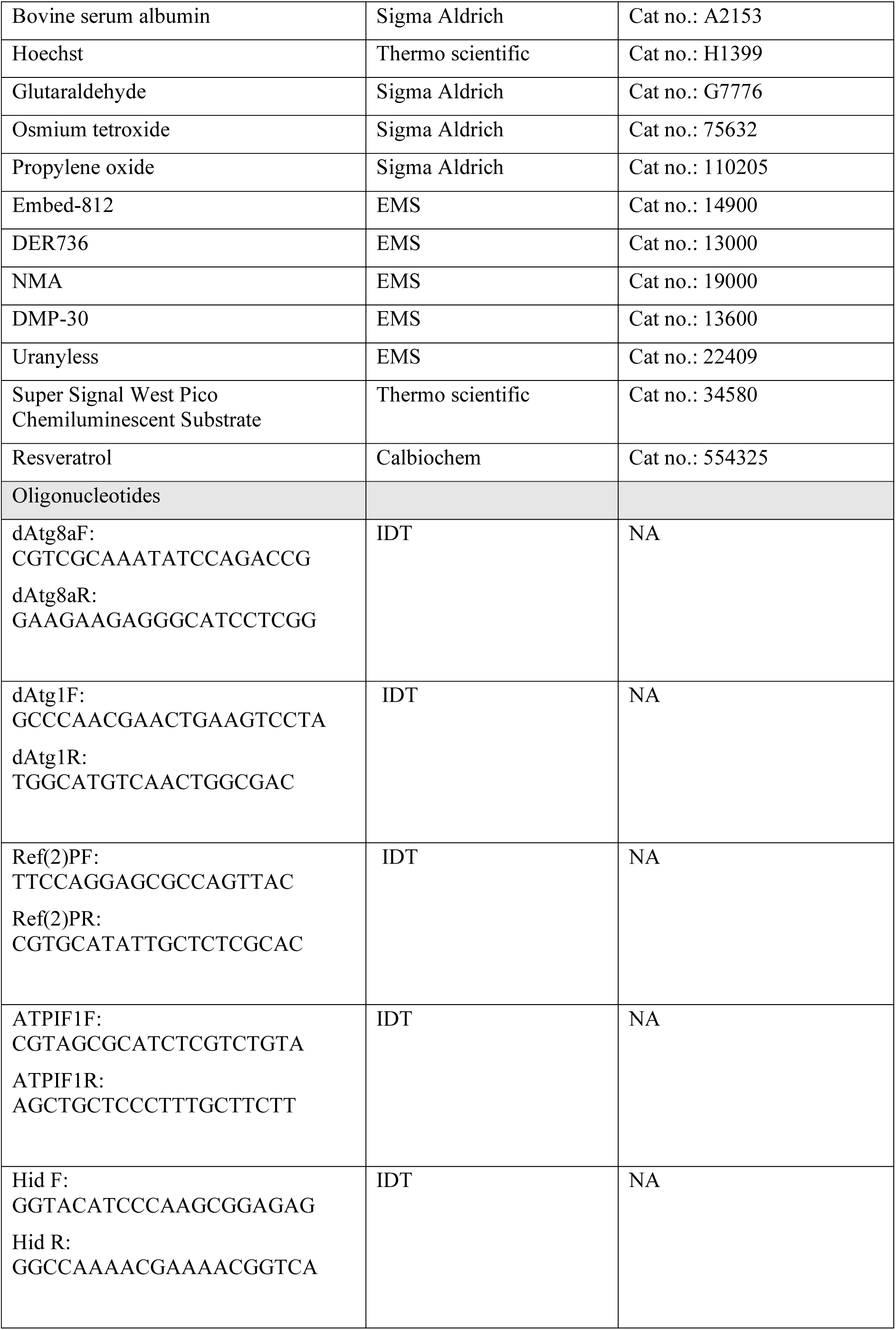
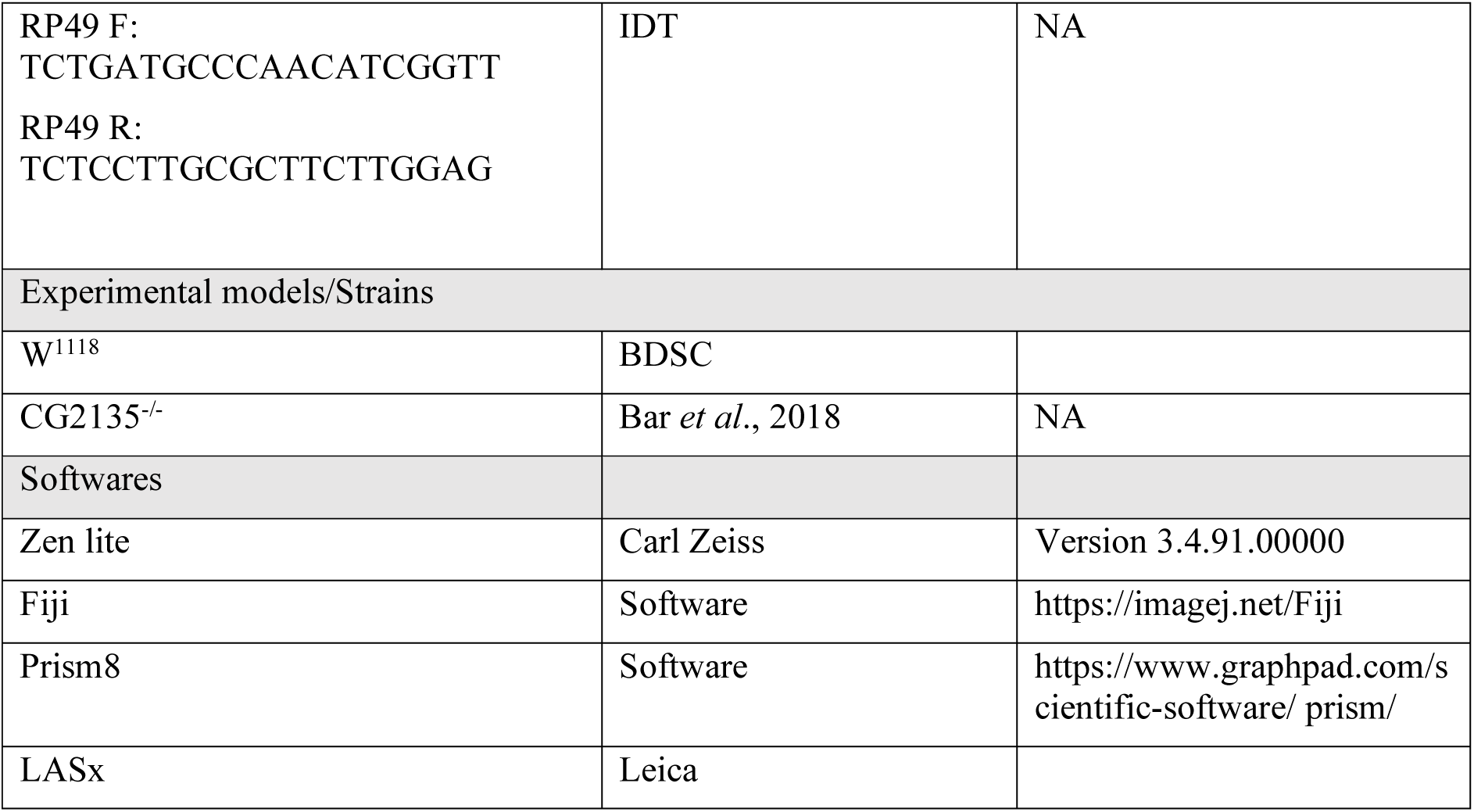

## EXPERIMENTAL MODEL AND METHOD DETAILS

### Drosophila strains and maintenance

W^1118^ flies were obtained from the Bloomington *Drosophila* Stock Centre at Indiana University, USA. The CG2135^-/-^ fly used in this study was reported earlier from our lab by Bar *et. al.* ^10^ and was maintained by routine outcrossing. W^1118^ and CG2135^-/-^ flies were maintained in controlled density at 25°C under a 12-hour day and night cycle. The standard cornmeal food (corn flour 75g/L, sugar 80g/L, agar 10g/L, dry yeast 15g/L, and malt extract 30g/L) media was used for raising the flies.

For age-matched flies, flies were allowed to lay eggs after synchronization for 24 hours in a cage setup. Then the freshly hatched larvae were collected in equal numbers in food vials. After 9-10 days the newly eclosed flies were collected and used for respective in the experiments. For all the experiments only male flies were used.

Resveratrol dissolved in 100% ethanol (Merck) was added to liquid cornmeal food at 400 µM concentration as previously reported ^10^. Flies were kept in continuous exposure of resveratrol starting at day 4 after eclosion till the day of experiment. Resveratrol containing food vials were replaced with fresh vials every alternative day.

### Starvation assay

Flies from W^1118^ and CG2135^-/-^ of 4 days, 15 days, and 30 days old were assayed for their survivability in no food condition. On the day of the experiment, flies in a cohort of 10 were kept in vials including a tissue paper soaked in distilled water to mimic no food condition and maintained under a constant temperature of 25°C with 12 hours dark and light cycle. The number of surviving flies monitored from their mobility was considered alive or dead and recorded every 12 hours. More than 130 flies were analysed for the starvation assay, and the experiment was done independently three times. The percentage of survivability was calculated from the total number of flies and live flies at each time point. To calculate mean survivability average number of flies that survived was determined ^89^.

### Western blot

For western blot, 5-10 fly heads were homogenized in 1x Laemmli’s (for 1ml 2X 0.5M Tris-Cl pH 6.8 0.312ml, SDS 0.05g, glycerol 0.25ml, 1M DTT 50µl) buffer supplemented with 1x protease inhibitor cocktail and boiled at 100°C for 5 min. An equal volume of the fly lysate was subjected to SDS-PAGE. Gels were transferred for 1 hour and blocked in 5% skim milk in 0.025% TBST. The primary antibody incubation was done at 4°C overnight in respective dilutions of antibodies. The secondary antibodies were incubated after washing for 1.5 hours. After final washing, the blots were developed using Super Signal West Pico Chemiluminescent Substrate and captured using the ChemiDoc imaging system (Biorad). α-Tubulin has been used as a loading control. Densitometric analysis of protein bands was quantified using ImageJ software.

### Isolation of mitochondria-rich fraction from fly head

Differential centrifugation was performed to isolate mitochondria from the fly head as described previously ^90^. Briefly, 50 fly heads were crushed gently with a mortar pestle (25 strokes) in 1 ml lysis buffer containing 250 mM sucrose, 0.1 mM EDTA and 2 mM EGTA. Homogenate was centrifuged at 350g for 3 min to eliminate the tissue debris. The supernatant was centrifuged twice at 1000g to eliminate the nuclear fraction. The mitochondria containing supernatant were centrifuged at 10000g for 10 min, the pellet was lysed in 1X Laemmeli’s buffer and western blot was performed. All the steps were performed in cold. Fractions were checked for the presence of mitochondrial and cytosolic proteins determined by immunoblotting with ATP5A and GAPDH, respectively.

### RNA isolation and qRT-PCR

Total RNA was isolated from 10-20 heads of W^1118^ and CG2135^-/-^ flies using TRIzol reagent following the manufacturer’s instruction. 1 µg of RNA was used for DNase treatment and DNA-free RNA was reverse transcribed using a cDNA synthesis kit. Transcript levels of different genes were quantified using this cDNA as the template. Quantitative real-time PCR (qRT-PCR) was performed with SYBR Green PCR reaction mix (BioRad) using the Biorad real-time PCR system. Target gene expression was normalized to RP49 as the reference gene (endogenous control). The results of quantitative real-time PCR were analysed by the comparative threshold cycle (CT) method ^91^.

### Immunostaining and imaging

Adult brains from W^1118^ and CG2135^-/-^ were dissected in S2 media supplemented with 10% FBS, 100 U/ml penicillin and 100 μg/ml streptomycin. The brains were fixed in 4% paraformaldehyde (PFA) for 30 minutes at room temperature. After fixation, the samples were washed in 0.3% PT (1x PBS + 0.3% triton X-100) and blocked with PBT x GS (0.3% PT+0.5% BSA+ 5% Normal goat serum) for 2 hours. The brains were incubated in primary antibodies at 4°C for 36 hours. Following incubation, the brains were washed 5x with 0.3% PT for 15 minutes under gentle shaking conditions and incubated in secondary antibody for 12 hours at 4°C. The washing steps were repeated and counter-stained with DAPI (1μg/ml). After the final wash for 15 minutes the brains were stored in Vectashield until use in the cold. The mounted brains were imaged in Apotome.2 microscope (Carl Zeiss) or Leica SP8 confocal microscope. The image processing was done using the ZEN blue software from Carl Zeiss and for quantification, ZEN blue software and ImageJ were used. For mean intensity measurement, 15-20 stacks from the Z planes were merged as MIP (maximum intensity projection) and the mean was taken from the multiple ROIs for each field using ZEN blue.

### MitoTracker, LysoTracker, and JC-I staining

MitoTracker staining for age-matched flies was done after dissecting the brain in S2 media ^10^. The samples were incubated in 500 nM MitoTracker for 1 hour and Hoechst (2μg/ml) with gentle shaking. After incubation 2 quick washes were given with S2 media. The brains were mounted in media and imaged immediately in Apotome.2 (Carl Zeiss). The intensity of the MitoTracker per unit (1000 x 1000 pixel unit area) area and mitochondrial size were measured using ImageJ software.

For MitoTracker and LysoTracker co-staining, brains were dissected in S2 media. Following dissection live brains were incubated with 500 nM MitoTracker in S2 media for 30 minutes with Hoechst (2μg/ml) hour at 25°C with gentle shaking in between. After 30 minutes, fresh Mitotracker along with 100 nM LysoTracker green and Hoechst was added and incubated for further 30 minutes. After incubation, 2 quick washes were given to remove the excess stain, followed by mounting in S2. The stained brains were immediately imaged with Apotome.2 40X objective. Colocalization between MitoTracker and LysoTracker was analysed using ImageJ. Single stacks of Z plane were taken and analysed using the JaCoP plugin.

For JC1 staining the brains were dissected in S2 media supplemented with 10% FBS. Dissected brains were incubated with 2 µM JC1 in S2+10% FBS for 45 mins. After incubation, the stain was discarded and washed quickly with S2+10% FBS. Washed brains were mounted in S2+10% FBS and imaged immediately using Apotome.2 (Carl Zeiss). The red-to-green ratio was calculated from individual stacks analysed in ImageJ.

### TUNEL staining

To determine the mechanism of cell death in W^1118^ and CG2135^-/-^ TUNEL assay was performed ^92^. The brains were dissected in PBS and fixed in 4% PFA in 0.1% triton for 30mins. Then the samples were washed 2 times each in 0.1% PT and 0.5% PT for 3 mins. The brains were permeabilized in sodium citrate at 65°C for 30 minutes followed by three washes in 0.5% PT. The samples were incubated with the TUNEL mix at 37°C for 3 hours following the manufacturer’s instructions. After three washes with 0.1% PT and counter-staining with DAPI, the brains were stored in Vectashield and imaged the next day in Apotome.2 (Carl Zeiss). Images acquired were processed in ZEN. 15-20 Z-stacks were taken for MIP and after threshold set for the samples count of TUNEL positive puncta was determined using ImageJ. The count has been represented per brain.

For co-staining with TH (tyrosine hydroxylase), the dissected brain was first immunostained with anti-TH antibody following the above-mentioned methodology used for the TUNEL staining protocol. The brains were mounted in Vectashield and imaged in 63x oil objective Leica SP8 confocal. TUNEL-positive dopaminergic neurons were analysed and manually counted.

### Lipofuscin imaging

Lipofuscin imaging was performed following previously described protocol ^93^. Briefly, brains from age-matched flies were dissected in cold phosphate buffer and kept on ice for 20 minutes. Then, they were immediately imaged in Apotome.2 (Carl Zeiss) using the FITC channel (Autofluorescence). The images were processed in ZEN blue. To count the number of autofluorescence bodies, MIP of 15 stacks was taken and the threshold was kept the same for all the samples and analysed in ImageJ.

### Tissue preparation and Electron microscopy (TEM)

Briefly, the dissected brains were fixed in 2.5% glutaraldehyde overnight, followed by post-fixation in 1% osmium tetroxide for 6h. The fixed tissues were dehydrated in a gradient of ethanol (30 to 100%) and finally in propylene oxide. Dehydrated tissues were infiltrated with resin (Embed-812 2ml, DER736 8ml, NMA 8.9ml and DMP-30 0.28ml) overnight. The next day, individual samples were embedded in capsule moulds and kept at 60°C for 48 hours to facilitate hardening. The samples were cut into ultra- thin using Leica Ultramicrotome. Sections of 70nm were negatively stained in uranyless. The images were acquired in Jeol JEM 2100Plus at 200 keV. The samples were analysed using ImageJ to determine the size and number of mitochondria and multilamellar bodies.

### ATP quantification

Total ATP was quantified using an ATP determination kit. For each set of measurement, 10-15 fly heads were homogenized in 1X PBS supplemented with 1X PIC in cold. Supernatant collected by centrifuging the homogenate at 20000g for 20 min was used to measure ATP. Total ATP quantified following manufacturer protocol using a standard curve generated with known concentrations of ATP. Total ATP content was normalized per µg of total protein.

### Statistical analysis

All the experiments were done at least three times independently. All statistical analysis were done using GraphPad Prism. *p*-values ≤0.05 were considered statistically significant as indicated by asterisks: \**P*≤0.05, \*\**P*≤0.01, and \*\*\**P*≤0.001.

---

**Figure S1:**
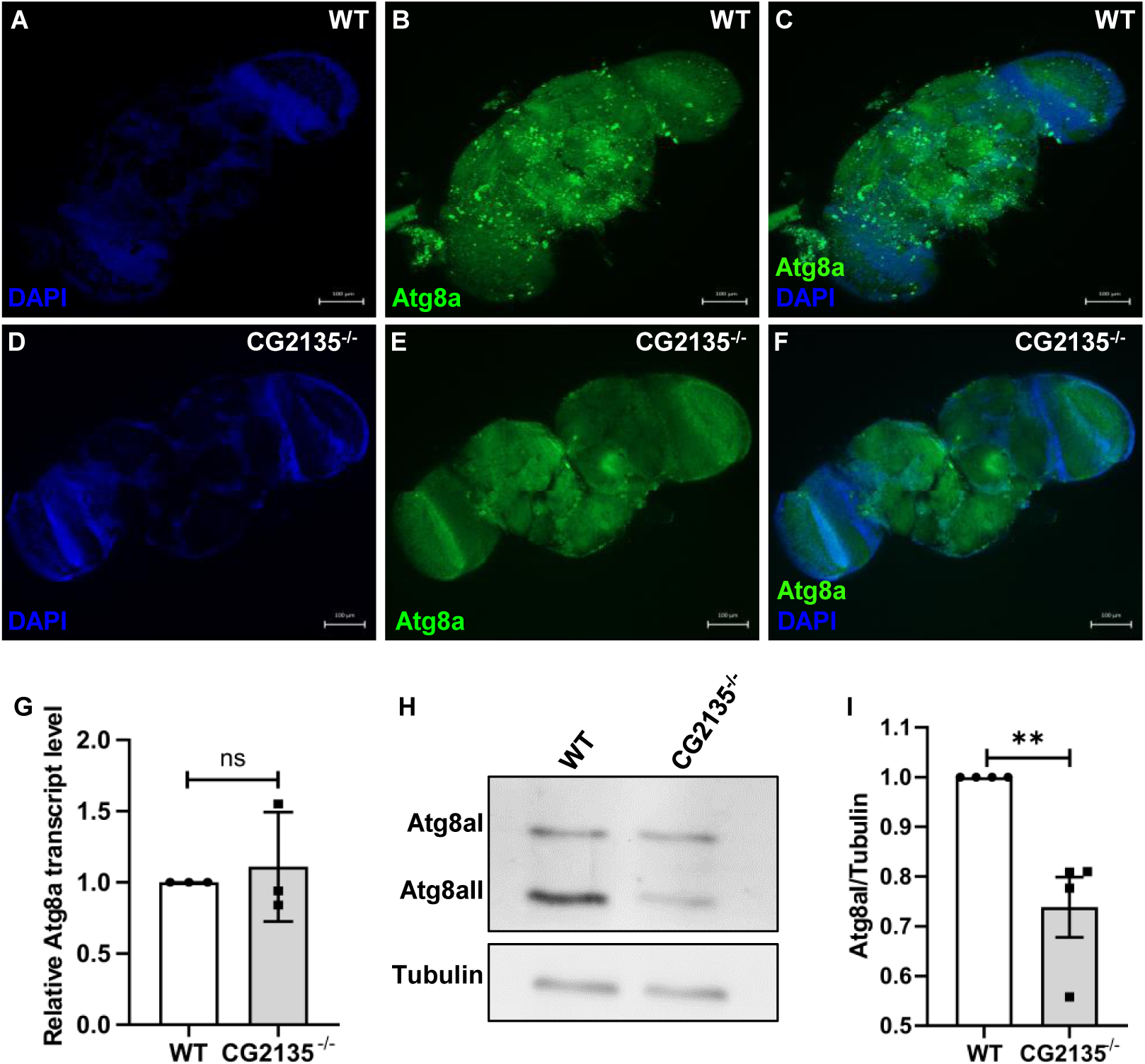
Reduced Atg8a in 30 days old CG2135^-/-^ fly brain, Related to Figure 2. (A-F) The whole brain of 30 days WT and CG2135^-/-^ was stained with anti-Atg8a (green) and counterstained with DAPI (blue), showing a low Atg8a signal in CG2135^-/-^ brain compared to WT. Images taken in Apotome.2 (Carl Zeiss) and the tiles were reconstructed using Zen software. Scale bar represents 100μm. (G) The bar graph represents the Atg8a transcript level in 30 days WT and CG2135^-/-^ determined by RT-qPCR normalised to RP49 as housekeeping gene control. (N=3) (H-I) Immunoblot of Atg8a for 30-day-old head lysate of WT and CG2135^-/-^. Relative quantification of Atg8a-I/Tubulin intensity from immunoblot and tubulin kept as a loading control. (N=3) All experiments were done independently. Error bars represent the standard error of the mean (SEM). Asterisks represent a level of significance (\*\**P* ≤ 0.01, Student’s *t*-test).

**Figure S2:**
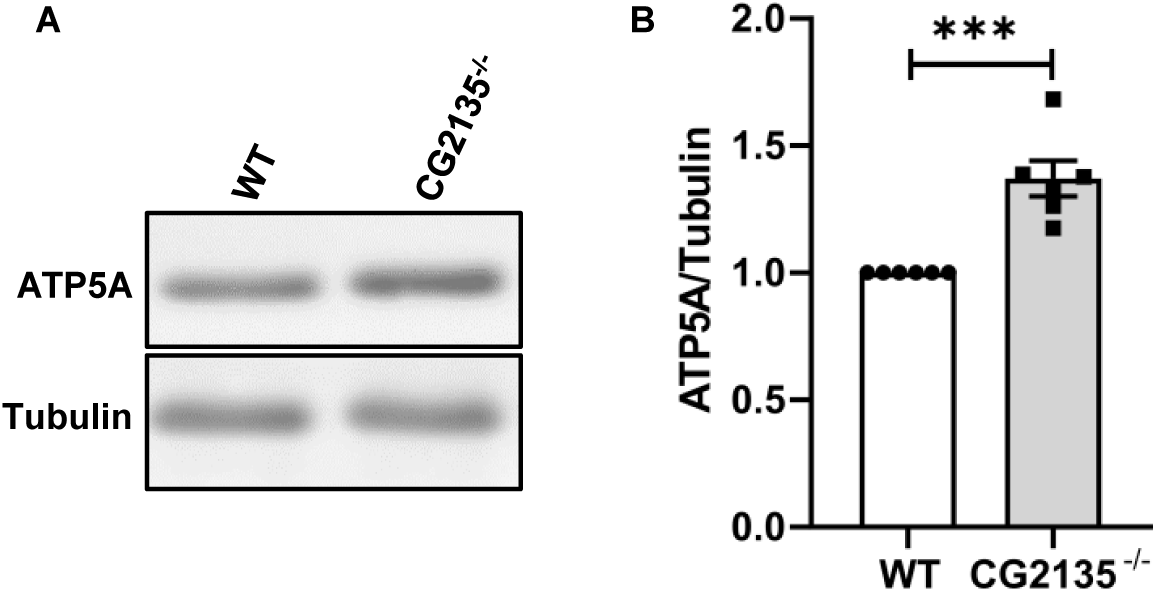
Western blot analysis of ATP5A protein, Related to Figure 3. (A-B) Western blot of mitochondrial protein ATP5A followed by quantification of the data shows an increased abundance of ATP5A in ≥30 days CG2135^-/-^ fly brain sample, suggesting increased accumulation of mitochondria. (N=6) All experiments were done independently. Error bars represent the standard error of mean (SEM). Asterisks represent a level of significance (\*\*\**P* ≤ 0.001, Student’s *t*-test).

**Figure S3:**
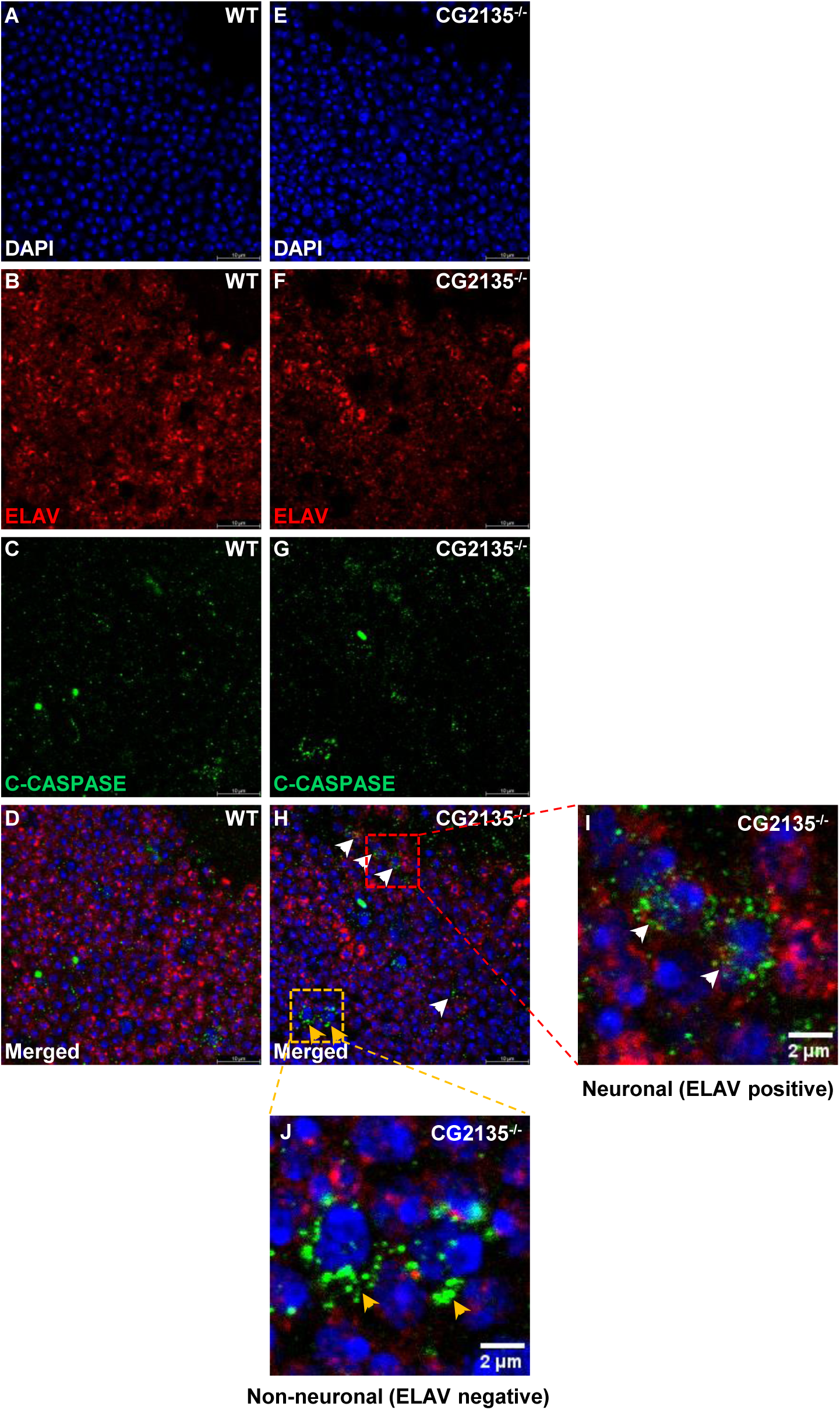
Accumulation of cleaved caspase3 positive cells in 45 days old CG2135^-/-^ fly brain, Related to Figure 6. (A-H) Co-immunostaining with anti-Cleaved caspase 3 and anti-ELAV showed ELAV positive cells (pan-neuronal) are Cleaved caspase 3 positive cells (white arrow). A few non-ELAV cells are also Cleaved-caspase 3 positive (yellow arrow). (I-J) Zoomed image showing ELAV positive (white arrow) pan-neuronal as well as non-Elav (yellow arrow) cells are cleaved caspase 3 positive suggesting a global apoptosis activation.

